# ANP32A and ANP32B are key factors in the Rev dependent CRM1 pathway for nuclear export of HIV-1 unspliced mRNA

**DOI:** 10.1101/559096

**Authors:** Yujie Wang, Haili Zhang, Lei Na, Cheng Du, Zhenyu Zhang, Yong-Hui Zheng, Xiaojun Wang

## Abstract

The nuclear export receptor CRM1 is an important regulator involved in the shuttling of various cellular and viral RNAs between the nucleus and the cytoplasm. HIV-1 Rev interacts with CRM1 in the late phase of HIV-1 replication to promote nuclear export of unspliced and single spliced HIV-1 transcripts. However, other cellular factors that are involved in the CRM1-dependent viral RNA nuclear export remain largely unknown. Here, we identified that ANP32A and ANP32B mediate the export of unspliced or partially spliced viral mRNA via interacting with Rev and CRM1. We found that double, but not single, knockout of ANP32A and ANP32B, significantly decreased the expression of gag protein. Reconstitution of either ANP32A or ANP32B restored the viral production equally. Disruption of both ANP32A and ANP32B expression led to a dramatic accumulation of unspliced viral mRNA in the nucleus. We further identified that ANP32A and ANP32B interact with both Rev and CRM1 to promote RNA transport. Our data strongly suggest that ANP32A and ANP32B play important role in the Rev-CRM1 pathway, which is essential for HIV-1 replication, and our findings provide a candidate therapeutic target for host defense against retroviral infection.

---

Human immunodeficiency virus type-1(HIV-1) is an enveloped, single strand positive RNA virus belonging to the lentivirus family. During HIV-1 replication, the 9 kb genome RNA is reverse transcribed to proviral DNA and then processed into RNA with specific splicing schemes to generate multiple species of mRNA: the unspliced 9 kb full-length genomic RNA, the 4 kb partially spliced mRNA and the 2 kb completely spliced mRNA (1). The mechanism for export of the unspliced and partially spliced viral RNAs is discrete from the pathway for transport of completely spliced mRNA. In the early phase of HIV-1 replication, the current consensus view of the completely spliced mRNA is presumably transported into the cytoplasm mediated by the TAP/NXF1 pathway to translate the regulatory proteins Rev, Nef, and Tat. The Rev protein of HIV-1, a 116 amino acid accessory protein, has a nuclear localization signal (NLS) recognized by importins, and also a leucine-rich nuclear export signal (NES) recognized by mammalian nuclear export factor Chromosomal Maintenance 1 (CRM1); it therefore shuttles between the nucleus and cytoplasm. Rev can specifically bind to the Rev response element (RRE) located in the *env* gene of the unspliced or partially spliced mRNA. In the late phase of HIV-1 replication, with Rev accumulation in the nucleus the unspliced or partially spliced mRNAs are exported to the cytoplasm via a Rev-CRM1 dependent export pathway to translate all structural viral proteins (2–10).

The key factor in the export of viral mRNA is the Rev-CRM1 complex. In the nucleus, the multimerized Rev recruits CRM1 through the Rev leucine-rich nuclear export signal (NES) located in the C-terminal domain to assemble the ribonucleoprotein complex with Ran-GTP(11–13), thus facilitating its exportation. However, a number of host factors (including Matrin-3, DDX1, DDX21, DDX3, DDX5, MOV10, Sam68, and CBP80) have been reported to interact with Rev and RRE and help viral mRNA export from the nucleus to the cytoplasm and mRNA translation (14–23). Most of these factors have not been identified as interacting with CRM1. So far there are only few proteins, such as DDX3 and Naf1, that are reported to interact with the Rev-CRM1 complex to facilitate viral mRNA export. Whether other cellular proteins are involved in the Rev-CRM1 complex and direct the viral RNA export from the nucleus to the cytoplasm remains largely unknown.

CRM1 is well known as an important receptor in the nuclear-cytoplasmic transportation of cellular RNA complexes and many viral RNAs, and functions by interacting with different cellular proteins through various mechanisms (24–27). In this study, we investigated the functions of two proteins that interact with CRM1, ANP32A and ANP32B, during the export of HIV-1 unspliced mRNA from the nucleus to the cytoplasm. The ANP32 family member, initially identified as a 32kDa highly conserved acidic (leucine-rich) nuclear phosphoprotein protein, is characterized by an N-terminal leucine rich repeat (LRR) domain and a C-terminal low-complexity acidic (LCAR) region (28). It has been suggested that the LRR domains of the ANP32 proteins interact with several proteins, including CRM1 (29), PP2Ac (30), Ataxin-1 (31), Histone H3-H4 (32), and Clip170 (33). The ANP32 proteins are also ascribed biochemical activities including chromatin modification and remodeling, apoptotic promotion (34), as well as protein phosphatase 2A (PP2A) and histone acetyltransferase (HAT) inhibition (35–37). It has been shown that ANP32A is an essential host partner co-opted to support influenza virus vRNP polymerase activity, and contributes to the influenza virus host range (38,39). Furthermore, it was reported that foamy virus nuclear RNA export is dependent on the HuR and CRM1 pathways in which ANP32A and ANP32B act as necessary cofactors (40). However, whether ANP32A/B involved in the Rev-dependent CRM1 pathways for HIV-1 unspliced RNA nuclear export is unknown.

Here, we created ANP32A and ANP32B knockout HEK293T cell lines to evaluate their functions in HIV-1 RNA export. Our work showed that ANP32A and ANP32B play similar roles and contribute to CRM1-dependent RNA export by interacting with both Rev and CRM1, therefore enhancing viral production and replication.

## RESULTS

### ANP32A and ANP32B are required for HIV-1 gag expression and viral production

ANP32A, encoded on chromosome 15, and ANP32B, encoded on chromosome 9, share 72.7% sequence identity and have similar protein structures, which consist of four acidic leucine-rich domain (LRR) repeats at the N terminal and a low-complexity acidic region (LCAR) located at the C terminal, bridged by a LRRCT domain (Fig. 1A). To determine whether human ANP32A and ANP32B could function as cellular co-factors in HIV-1 replication, we analyzed the effect of ANP32A and ANP32B gene knockout on HIV-1 gag expression and virus production. Firstly, two specific sgRNAs were used to generate ANP32A and/or ANP32B gene knocked out cell lines in HEK293T cells using CRISPR/Cas9 technology (Fig. 1B). The ANP32A single knockout cell line ∆ANP32A (AKO), ANP32B single knockout cell line ∆ANP32B (BKO), and the ANP32A/B double KO cell line (DKO) were confirmed by western blotting (Fig. 1C) and gene sequencing (data not shown).

**FIG 1.**
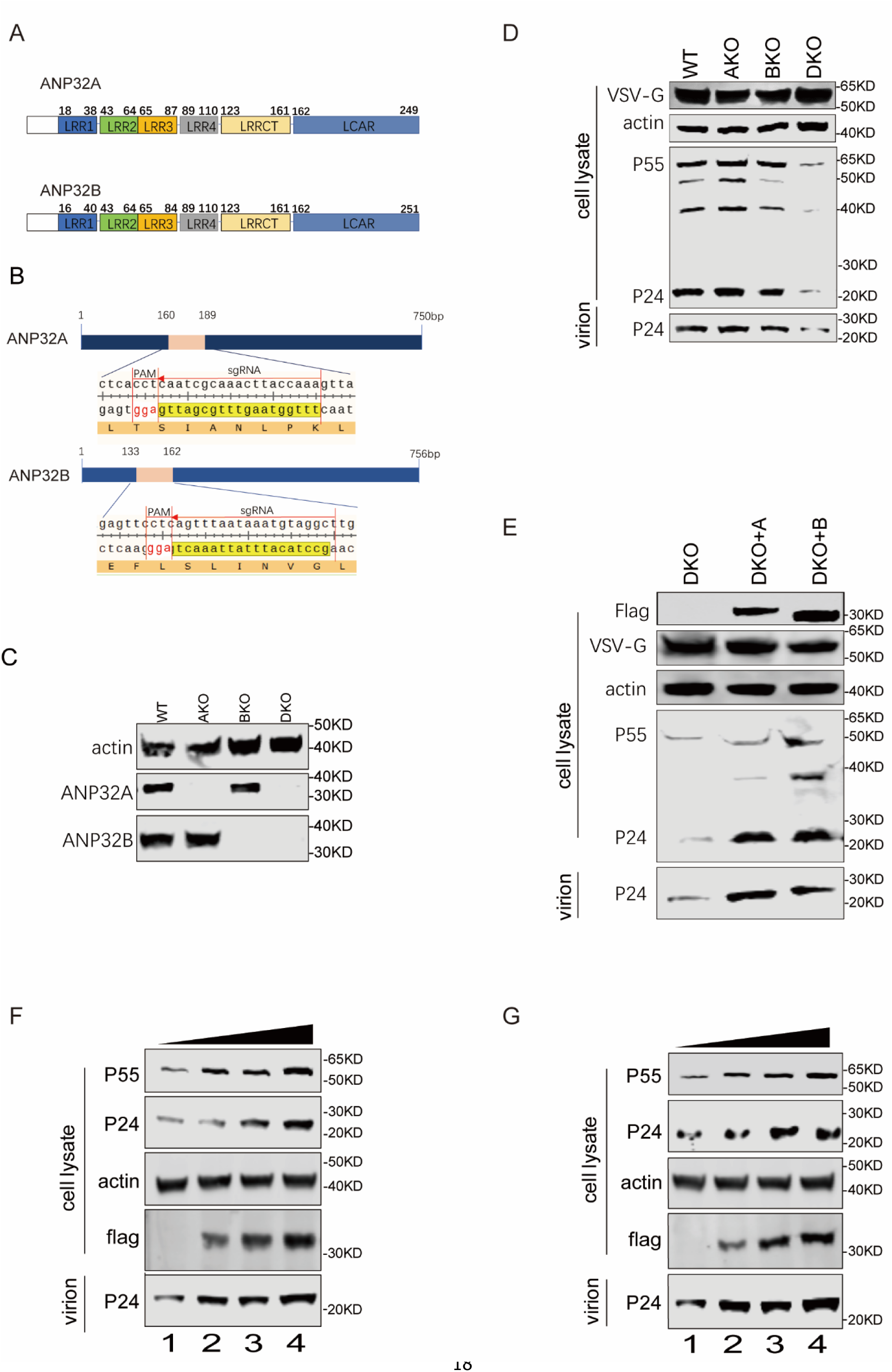
ANP32A and ANP32B are required for HIV-1 gag expression and viral production. (A) Schematic diagram of sequence analysis and secondary structure comparison of human ANP32A and ANP32B. (B) A schematic of ANP32A and ANP32B KO by CRISPR/Cas9 technology. The position of each sgRNA is depicted with yellow. (C) Verification of ANP32A, ANP32B, and actin expression by Western blotting in WT, AKO, BKO and DKO cells. The antibody for ANP32A and ANP32B were purchased from Abcam and actin antibody was purchased from sigma. (D) Gag expression in cell lysate and supernatants were identified by Western blotting. WT, AKO, BKO and DKO cells were transfected with pNL4-3-lucΔVifΔEnv and VSV-G-expressing plasmids. Virions were purified from cell culture supernatants by ultra-centrifugation 48 h post-transfection. Cells and virions were lysed and analyzed by Western blotting with the appropriate primary antibodies, including a mouse monoclonal anti-P24 antibody (Sino Biological), Rabbit Polyclonal VSV-G tag antibody (GeneTex), a mouse anti Flag antibody and mouse anti β-actin antibody (Sigma). (E) Rescue of p24 and p55 expression in the stable expressing ANP32A or ANP32B protein DKO cells. DKO, DKO+A or DKO+B were transfected with pNL4-3-lucΔVifΔEnv and VSV-G plasmids and HIV-1 production was quantified by measuring p24 and p55 expression 48 h later in cell lysates and cell culture supernatants. (F~G) Increasing amounts of human ANP32A (F) or human ANP32B (G) expression vectors, and virion protein expression in cells and culture supernatants were detected by Western blotting.

To determine the effect of ANP32A and ANP32B on HIV-1 production, we transfected pNL4-3-lucΔVifΔEnv and VSV-G plasmids into WT, AKO, BKO, and DKO cells and assessed the expression of viral Gag proteins p55 and p24. The results showed that single knockout of either ANP32A or ANP32B has no impact on HIV-1 gag expression, while a significant decrease in p24 expression in the cytosol and cell culture supernatant was observed in the DKO cells compared with WT, AKO and BKO cells (Fig. 1D). This result indicates that both ANP32A and ANP32B are functional and play similar role in HIV-1 gag expression and either is enough to support the virus protein expression. Furthermore, we confirmed that p24 expression was restored when ANP32A and ANP32B were reconstituted via ectopic stable expression in the DKO cells (Fig. 1E). In addition, when ANP32A or ANP32B-expressing plasmid was transfected into DKO cells, the p24 and p55 expression in the cell and cell culture supernatant both increased in a dose-dependent manner (Fig. 1F and G). All these results demonstrate that either ANP32A or ANP32B is required for HIV-1 gag expression in HEK293T cells.

### ANP32A and ANP32B enhance the export of unspliced and incompletely spliced transcripts from nucleus to cytoplasm

To clarify the mechanism by which ANP32A and ANP32B promote HIV-1 gag expression, the potential effect of ANP32A and ANP32B on viral reverse transcripts was investigated. HIV-1 pseudotyped virus was used to infect WT and DKO cells. Total DNA was extracted, and the viral early or late reverse transcripts were quantified by real-time PCR with specific probes at 2, 6 and 18 h post infection. There was no obvious effect on either the early or late reverse transcription products in DKO cells compared with WT (Fig. 2A and B).

**FIG 2.**
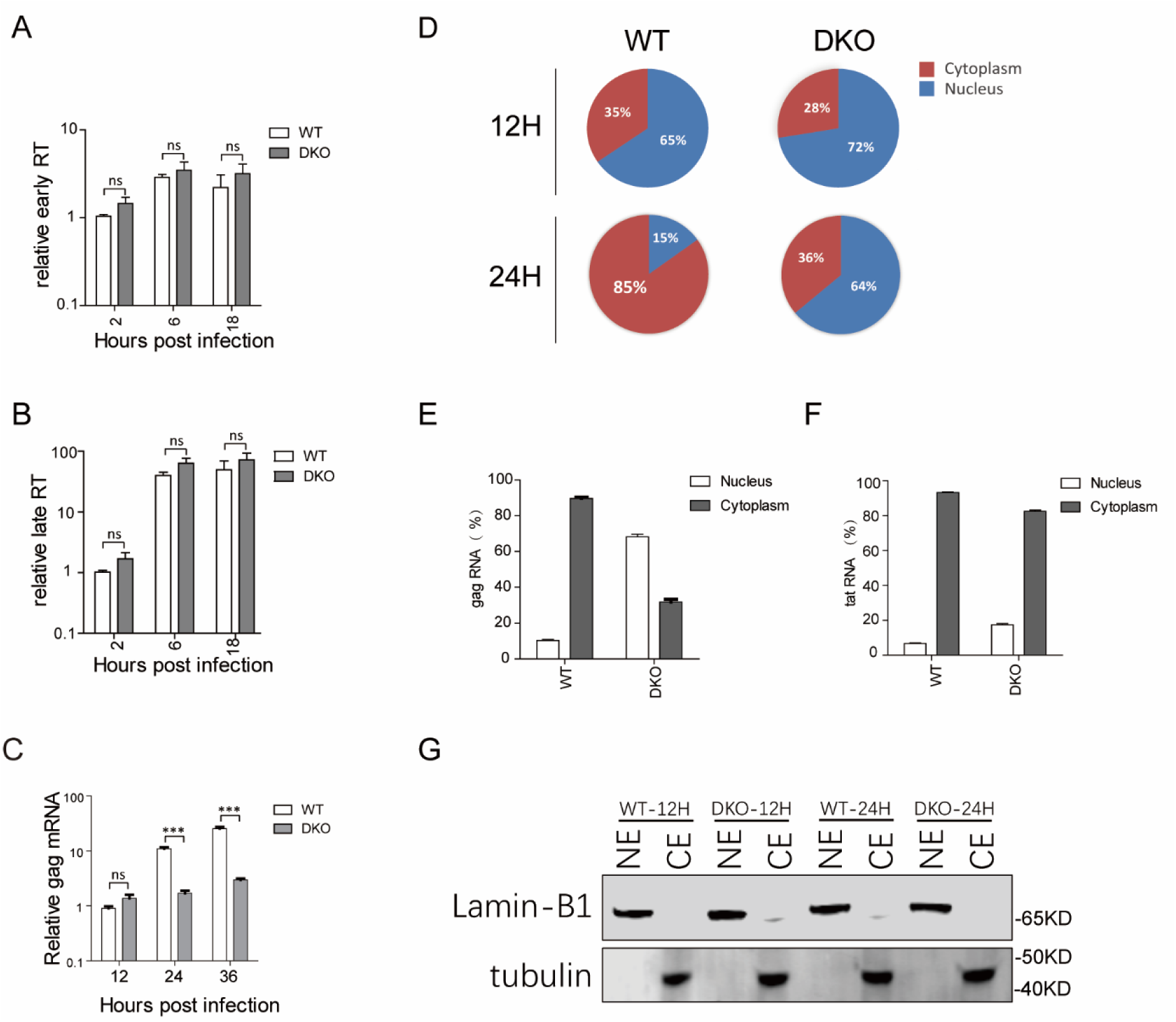
ANP32A and ANP32B enhance the export of unspliced mRNA from nucleus to cytoplasm. Both WT and DKO cells were infected with HIV-1 pseudotype virus. (A, B) Levels of accumulation of viral reverse transcripts. HIV-1 pseudotype virions were first treated with DNase and then used to infect WT and DKO cells. 2, 6, and 18 h later, cellular DNAs were extracted from these infected cells, and the viral early (A) and late reverse transcripts (B) were quantified by real-time PCR using specific primers and probes. Globin mRNA was measured as an endogenous control. (C) Total RNA was extracted from transfected cells and analyzed by real-time PCR using primers specific for unspliced mRNA. β-actin mRNA was measured as an endogenous control. (D) Cytoplasmic and nuclear RNAs isolated from WT and DKO cells at 12, 24 h post infection was analyzed by real-time PCR with primers specific for unspliced mRNA. (E) Cytoplasmic and nuclear unspliced *gag* RNAs distribution in the WT and DKO cells at the 24 h post infection was analyzed by real-time PCR with primers specific for gag mRNA. (F) Cytoplasmic and nuclear completely spliced *tat* RNAs distribution in the WT and DKO cells at the 24 h post infection was analyzed by real-time PCR with specific primers. The cytoplasm and nucleus relative ratios were then calculated and are presented. (G) Lamin-B1 and tubulin was used as nuclear or cytoplasmic control, respectively. Error bars represent standard deviation calculated from three experiments. ***p < 0.001, NS, not significant (p > 0.05).

We then tested the function of ANP32A and ANP32B in viral transcription and transport. HIV-1 pseudotyped virus was used to infect the WT or DKO cells and cellular RNA was collected to analyze the total *gag* mRNA. Our results showed that total *gag* RNAs in WT and DKO cells were similar at 12 h post infection, while at 24 and 36 h post infection, levels of viral *gag* mRNA in DKO cells were about 10~fold lower than in WT cells (Fig. 2C). This result suggests the existence of a block to viral RNA processing after RNA generation in DKO cells. ANP32A and ANP32B have been reported to bridge HuR and CRM1 to support FV RNA transport (40) and also to contribute to cellular RNA transport (29,41). We next examined whether ANP32A and ANP32B had any effect on HIV-1 unspliced mRNA nucleo-cytoplasmic shuttling. We fractioned the nucleus from the cytosol, and the cytoplasmic and nuclear distributions of HIV-1 unspliced mRNA in WT or DKO cells were analyzed by quantitative PCR at 12 h and 24 h post infection. At 12 h post infection, there is no obvious difference in the distribution of viral RNA between the WT and DKO cells, in that most of the viral RNA in both cell types was located in the nucleus. In contrast, at 24 h post infection, in the WT cells 85% of the RNA was located in the cytoplasm, indicating that the viral RNA was largely exported from the nucleus into the cytoplasm. However, in the DKO cells, most of the unspliced viral RNA had accumulated in the nucleus. As the result of the viral RNA retention in the nucleus, the ratio of transported viral unspliced mRNA vs total RNA in DKO cells at 24 h post infection was very low (Fig 2D). The transportation efficiency of viral *gag* mRNA from nucleus to cytoplasm was largely reduced in DKO cells compared with WT (Fig 2E). However, the completely spliced *tat* mRNA is mostly located in the cytoplasm at 24 post infection in both WT and DKO cells. Only slight alteration of the nuclear export of the *tat* mRNA was observed in DKO cells (Fig 2F). We used Lamin B1 and tubulin protein as nuclear and cytoplasmic protein controls respectively to assess the purity of the nuclear and cytoplasmic extracts (Fig. 2G). This result implied that ANP32A and ANP32B are required for the transport of unspliced viral mRNA from the nucleus to the cytoplasm. Absence of ANP32A and ANP32B could lead to the accumulation of gag mRNA in the nucleus and subsequently reduced gag mRNA production *in trans* (Fig 2C).

To better illustrate this phenomenon, the subcellular distribution of unspliced *gag* transcripts were visualized using fluorescence in-situ hybridization (RNA Scope) with RNA probes that specifically target *gag-pol* mRNA. Consistent with the above observations, at 24 and 36 h post infection, *gag-pol* mRNA in the WT cells is mostly located in the cytoplasm (Fig. 3A), whereas most of the *gag-pol* mRNAs were in the nucleus of the DKO cells (Fig. 3B), indicating a defect in the nuclear export of the unspliced transcripts in the absence of ANP32A and ANP32B. The distribution of the *gag-pol* mRNA observed using the DeltaVision OMX 3D structured illumination microscope (3D-SIM) (GE, USA) closely mirrored the confocal microscopy result. It was clearly showed that in the WT cells, the *gag-pol* mRNA RNA is mostly located in the cytoplasm (Fig. 3C), but in the DKO cells the unspliced mRNA was retained in the nucleus (Fig. 3D). These data strongly support that ANP32A and ANP32B contribute to the efficient export of unspliced viral *gag-pol* transcripts from the nucleus to the cytoplasm.

**FIG 3.**
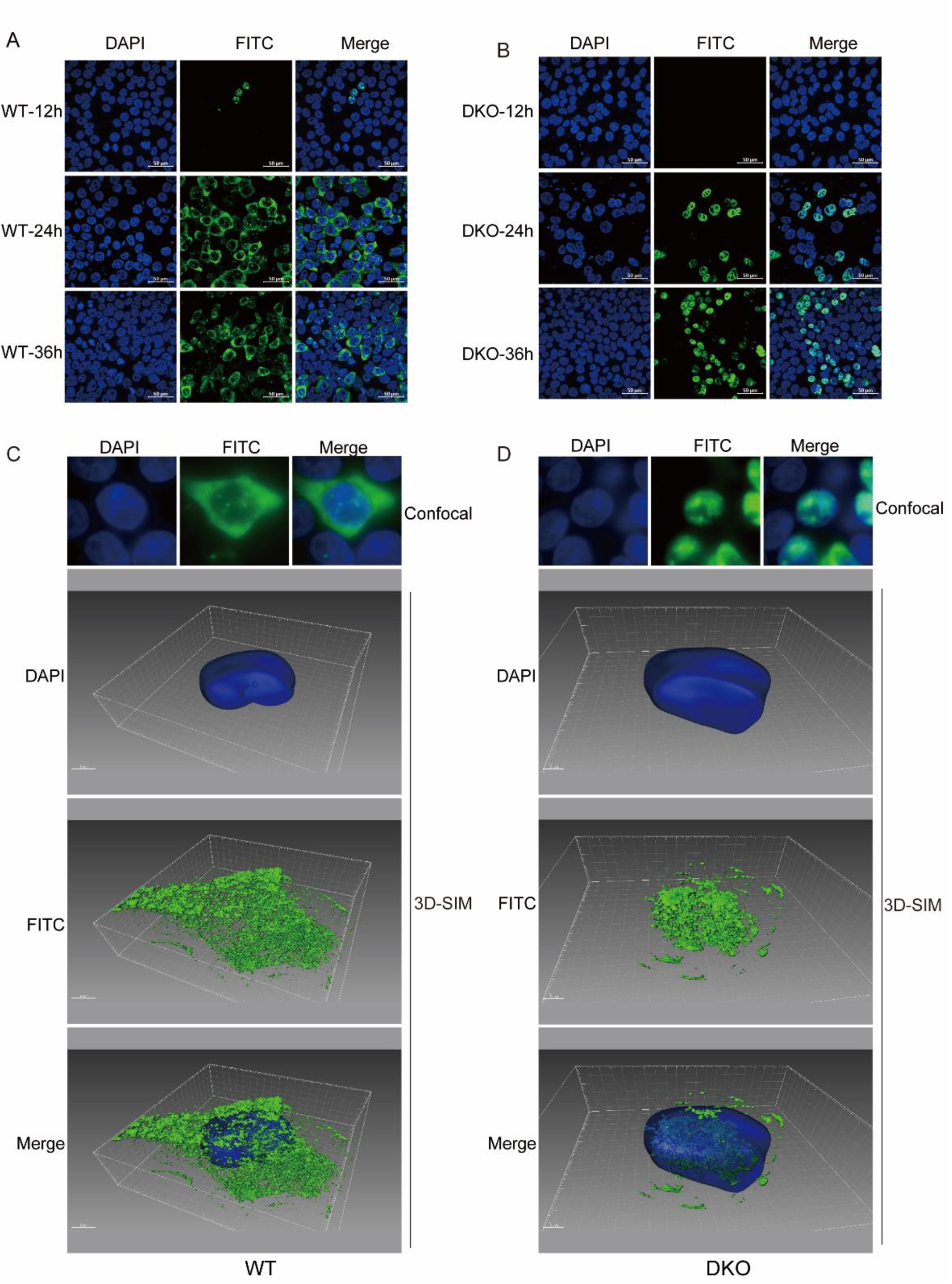
ANP32A and ANP32B double deletion reduces the levels of unspliced HIV-1 mRNA in the cytoplasm. (A) WT cells were infected with the HIV-1 pseudotype virus and cells were fixed, and stained with fluorescent *gag-pol*-specific RNA probes at 12, 24 and 36 h post infection. (B) DKO cells were infected with the HIV-1 pseudotype virus and cells were fixed and stained with fluorescent *gag-pol*-specific RNA probes at 12, 24 and 36 h post infection. DAPI staining was performed to visualize nuclei. Stained sections were captured and processed using fluorescence microscopy (ZEISS, LSM880, Germany). (C) WT cells were infected with the HIV-1 pseudotype virus for 24 h and cells were fixed, stained with fluorescent *gag-pol*-specific RNA probes. DAPI staining was performed to visualize nuclei. Stained sections were captured and processed using the Delta vision OMX SR imaging system (GE). (D) DKO cells were infected with the HIV-1 pseudotype virus for 24 h and cells were fixed, stained with fluorescent *gag-pol*-specific RNA probes. DAPI staining was performed to visualize nuclei. Stained sections were captured and processed using the Delta vision OMX SR imaging system (GE). The experiments were performed independently at least three times, with a representative experiment being shown.

### ANP32A and ANP32B interact with Rev

Rev is the key viral protein mediates unspliced viral RNA export from nucleus to cytoplasm. To investigate whether ANP32A and ANP32B interact with Rev, Rev-HA was transfected into HEK293T cells with ANP32A-Flag or ANP32B-Flag and the formaldehyde cross-linking immunoprecipitates (42,43) were analyzed by Western blotting in 24 h. The result showed that both ANP32A and ANP32B co-immunoprecipitate with Rev protein (Fig. 4A). We then analyzed the subcelluar location of Rev-HA and ANP32A-Flag or ANP32B-Flag in HeLa cells using confocal imaging. We observed that the ANP32A and ANP32B proteins were mainly found in the nucleus of the transfected cells. As Rev is a nuclear-cytoplasmic shuttling protein, we would expect to see it both free and co-localized with the proteins it shuttles. Our results suggest that a fraction of the Rev protein does indeed co-localize with ANP32A or ANP32B in the nuclei of the HeLa cells (Fig 4B).

**FIG 4.**
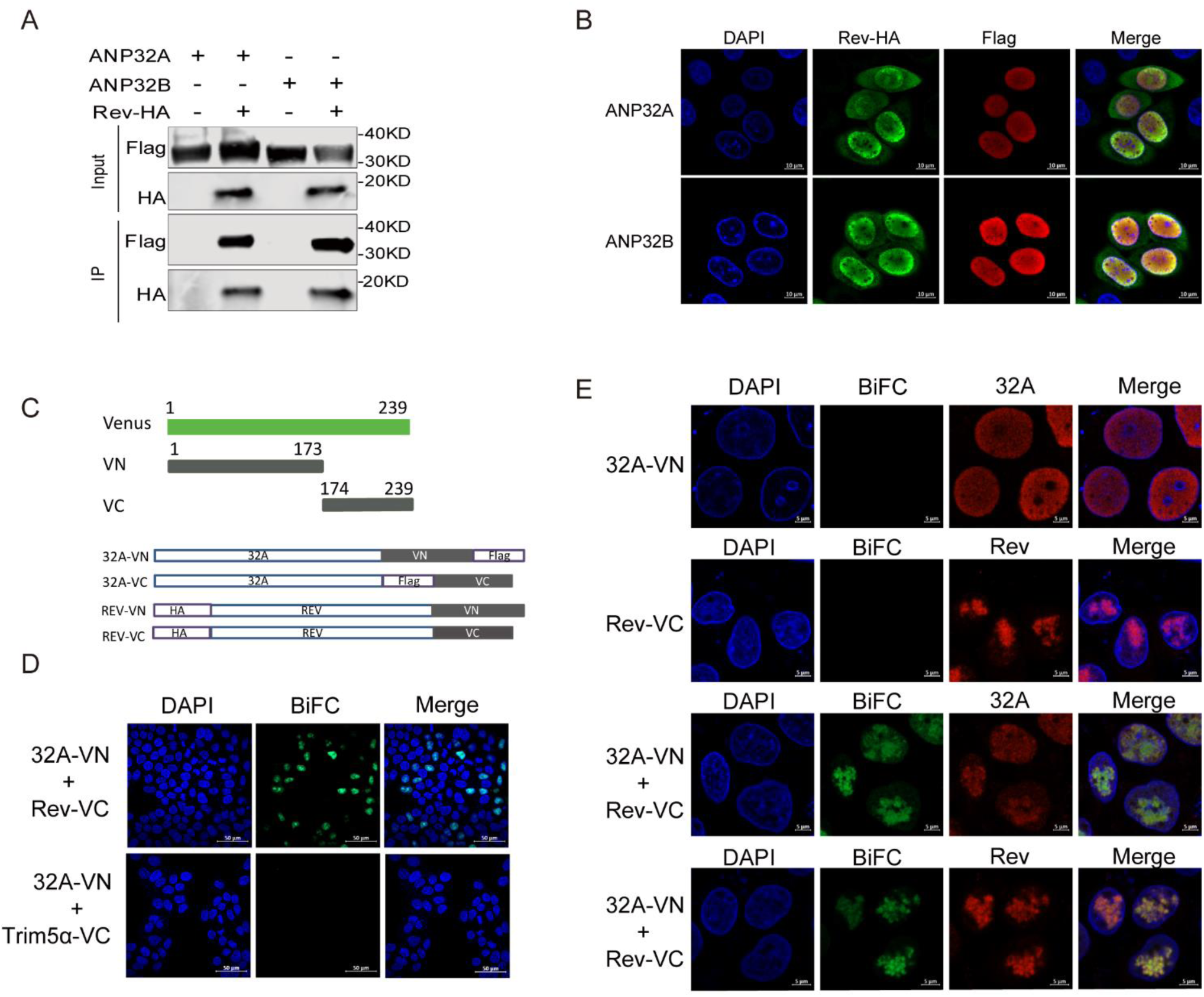
ANP32A and ANP32B interact with Rev. (A) Detection of ANP32A and ANP32B interactions with Rev. HEK293T cells were co-transfected with plasmids expressing ANP32A-Flag or ANP32B–Flag alone or in the presence of Rev-HA vector, the whole-cell lysates were prepared, and immunoprecipitations were performed with anti-HA beads and analyzed by Western blotting. (B) HeLa cells were transfected with 1 μg ANP32A or ANP32B with 1 μg Rev-HA expressing plasmids, and immunofluorescence stained to visualize the nucleus (DAPI), ANP32A or ANP32B-Flag (red) and Rev-HA (green). (C) A schematic of the BiFC fusion proteins is presented. Venus is divided into an N-terminal region from residues 2 to 173 (VN) and a C-terminal region from residues 174 to 239 (VC). VN and VC were fused to the C-terminus of the target proteins. (D) ANP32A-VN-Flag and HA-Rev-VC or VC-Flag-trim5α were co-transfected into Hela cells and the BiFC green fluorescent signals were visualized by confocal microscopy. (E) ANP32A-VN-Flag and HA-Rev-VC were expressed individually or together in Hela cells and ANP32A-VN-Flag and HA-Rev-VC fusion proteins were stained with a rabbit anti-Flag and anti-HA monoclonal antibody at 1:500 followed by an Alexa-Fluor-647-conjuated goat anti-rabbit antibody at 1:1000 dilution. The BiFC green fluorescent signals were visualized by confocal microscopy.

To further confirm the interaction between ANP32 proteins and Rev, a bimolecular fluorescence complementation (BiFC) assay was used. In our research, the function of ANP32B is similar to ANP32A, so in the BiFC assay we just detected the interaction between ANP32A and Rev. This assay enables direct visualization of protein interaction and the subcellular location in living cells. The BiFC assay is based on the finding that two non-fluorescent fragments of a fluorescent protein can associate to produce a significantly brighter fluorescent signal, and that a fluorescent signal can still be obtained if the association of the fragments is adjacent, for example if they are fused to specifically interacting partners. N-terminal residues 1-173 (VN) and C-terminal residues 174-239 (VC) of Venus were fused to the C-terminus of ANP32A and Rev protein, respectively (Fig. 4C). Firstly, we confirm that the Rev-VC was functional equally as wild type Rev in a RRE dependent reporter system (data not shown). When ANP32A-VN and Rev-VC were co-transfected into Hela cells, the Venus signal mostly located in the nucleus, while the human trim5α-VC plasmid was used as a negative control (Fig. 4D). The Venus signal was observed in the nucleus only when ANP32A-VN and Rev-VC were co-transfected compared with single transfection of ANP32A-VN or Rev-VC (no Venus signal) (Fig. 4E). Thus, together, these data indicate that both ANP32A and ANP32B can specifically interact with Rev in the nucleus of Hela cells.

### ANP32A and ANP32B play key role in the CRM1 dependent RNA export pathway

HIV-1 Rev/RRE dependent mRNA transport relies on the CRM1 export pathway. We next investigated whether ANP32A and ANP32B interact with CRM1. To do this, we overexpressed ANP32A-Flag or ANP32B-Flag, either with or without a CRM1 express vector (CRM1-HA), into HEK293T cells. Formaldehyde cross-linking immunoprecipitants with HA beads were performed. The result showed that both ANP32A and ANP32B co-immunoprecipitated with CRM1 (Fig. 5A). By confocal imaging analysis, we found that either ANP32A or ANP32B protein co-localized with CRM1 in the nucleus and perinuclear region (Fig. 5B). To further confirm the interaction between CRM1 and ANP32A, we performed a BiFC assay by using CRM1-VN and ANP32A-VC fusion proteins. We observed that ANP32A interacted with CRM1 at the nuclear periphery at 24 h post transfection when CRM1-VN and ANP32A-VC were co-transfected into Hela cells (Fig. 5C). When CRM1-VN or ANP32A-VC was transfected individually into Hela cells, the CRM1-VN was mostly located in the nuclear periphery and nucleus and ANP32A-VN protein was mostly located in the nucleus (Fig. 5C). These results confirm that ANP32A and ANP32B interact with CRM1 proteins.

**FIG 5.**
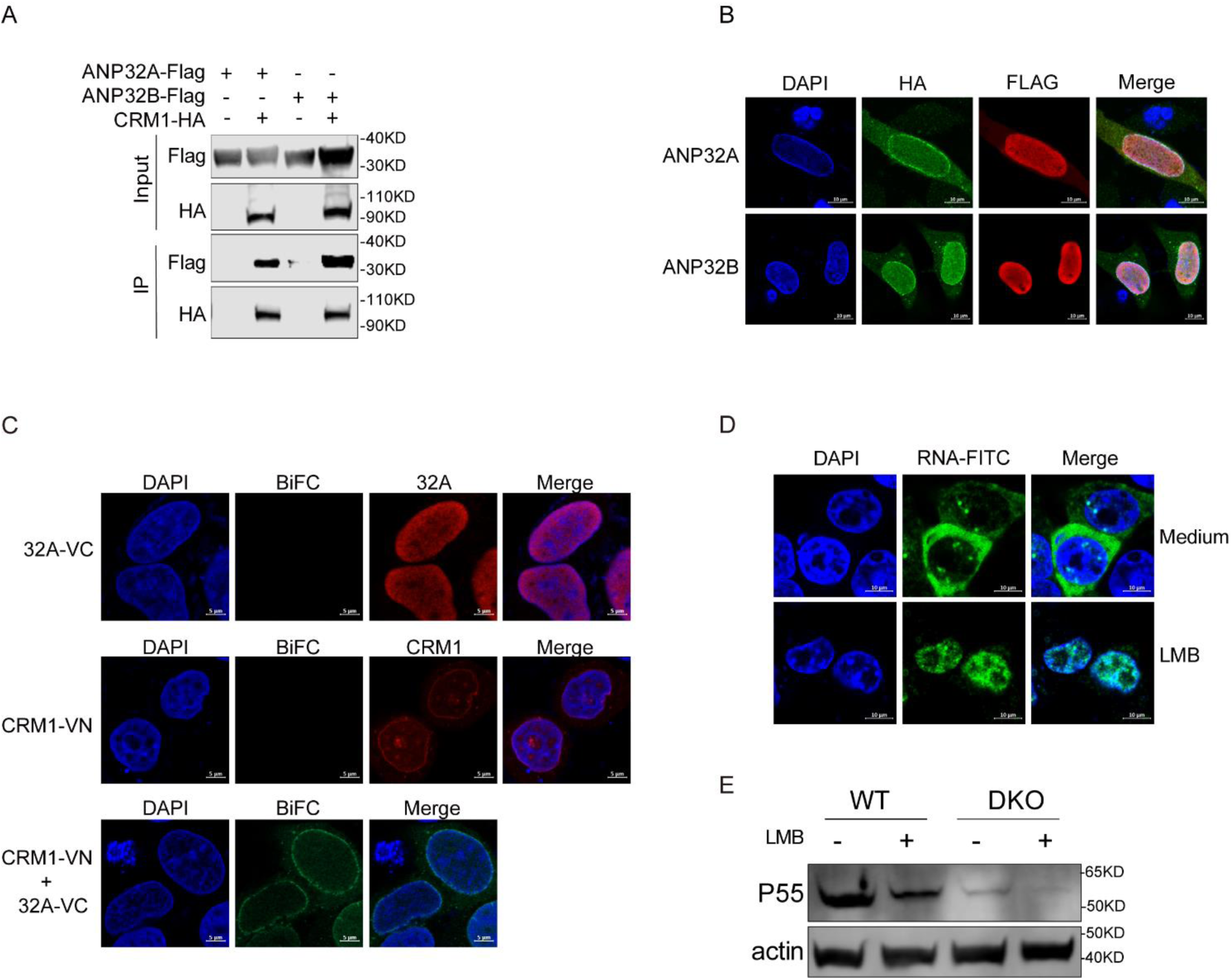
ANP32A and ANP32B play key role in the CRM1 dependent RNA export pathway. (A) Detection of ANP32A interaction with CRM1. HEK293T cells were co-transfected with plasmids expressing ANP32A-Flag alone or in the presence of the CRM1-HA vector, the whole-cell lysates were prepared, and immunoprecipitations were performed with anti-HA beads and analyzed by Western blotting. (B) HeLa cells were co-transfected with CRM1-HA and ANP32A-Flag or ANP32B-Flag expressing plasmids and immunofluorescence stained to visualize the nucleus (DAPI), ANP32A-Flag or ANP32B-Flag (red) and CRM1-HA (green). (C) CRM1-VN and ANP32A-VC were co-transfected into Hela cells and the BiFC green fluorescent signals were visualized by confocal microscopy. (D) The HEK293T cells were infected with HIV-1 psuedotype virus for 12 h and either medium or LMB (20 nM) (Beyotime) were added for additional 12 h. HIV-1 *gag* mRNA location was analyzed with a specific probe using an RNA Scope technology. (E) WT and DKO cells were transfected with pNL4-3-lucΔVifΔEnv plasmids for 12 h and either medium or LMB (20 nM) were added for additional 12 h. The HIV-1 production was quantified by measuring p55 expression.

Leptomycin B (LMB), a specific inhibitor that covalently binds to cys528 of CRM1 was used to block CRM1 mediated export from the nucleus (44). We found that HIV-1 unspliced mRNA transportation from the nucleus to the cytoplasm was significantly inhibited with LMB treatment (Fig. 5D). To further clarify the ability of ANP32A and ANP32B in the CRM1 dependent RNA export pathway, pNL4-3-lucΔVifΔEnv were used to transfect both WT and DKO cells either with or without LMB treatment. We found that in the WT cells, LMB treatment significantly decreased the efficiency of HIV-1 gag expression. Interestingly, depletion of ANP32A and ANP32B resulted in worse p55 expression, although the LMB treatment caused further reduction of gag expression in DKO cells (Fig. 5E). Considering the interactions between ANP32A/B and Rev/CRM1 together, we propose that ANP32A and ANP32B are required for the Rev/CRM1 dependent viral RNA export.

In the nucleus, Rev binds to the RRE and interacts with CRM1, facilitating nuclear export of viral RNAs. In the above result, we confirmed that ANP32A/B interact with Rev and CRM1 protein. To determine the functional relationship between ANP32A/B and Rev-CRM1 nuclear export pathway, we investigated whether the ANP32A/B deletion influence the interaction between Rev and CRM1. 293T WT and DKO cells were co-transfected with FLAG-tagged Rev with or without HA-tagged CRM1, co-IP experiments were performed. Cell lysates were formaldehyde cross-linking immunoprecipitated with an anti-HA beads, followed by western blotting. The result showed that either in the WT or DKO cells Rev protein interacts with CRM1 (Fig. 6A). The result was confirmed by the BiFC assay (Fig. 6B). This data demonstrated that deletion of ANP32A/B impaired CRM1/Rev/RRE dependent RNA transport without disruption of the interaction between Rev and CRM1. Together, we conclude that ANP32A and ANP32B are key cofactors of Rev, and they interact with CRM1 to mediate export of unspliced or partially spliced viral RNA from nucleus to cytoplasm.

**FIG 6.**
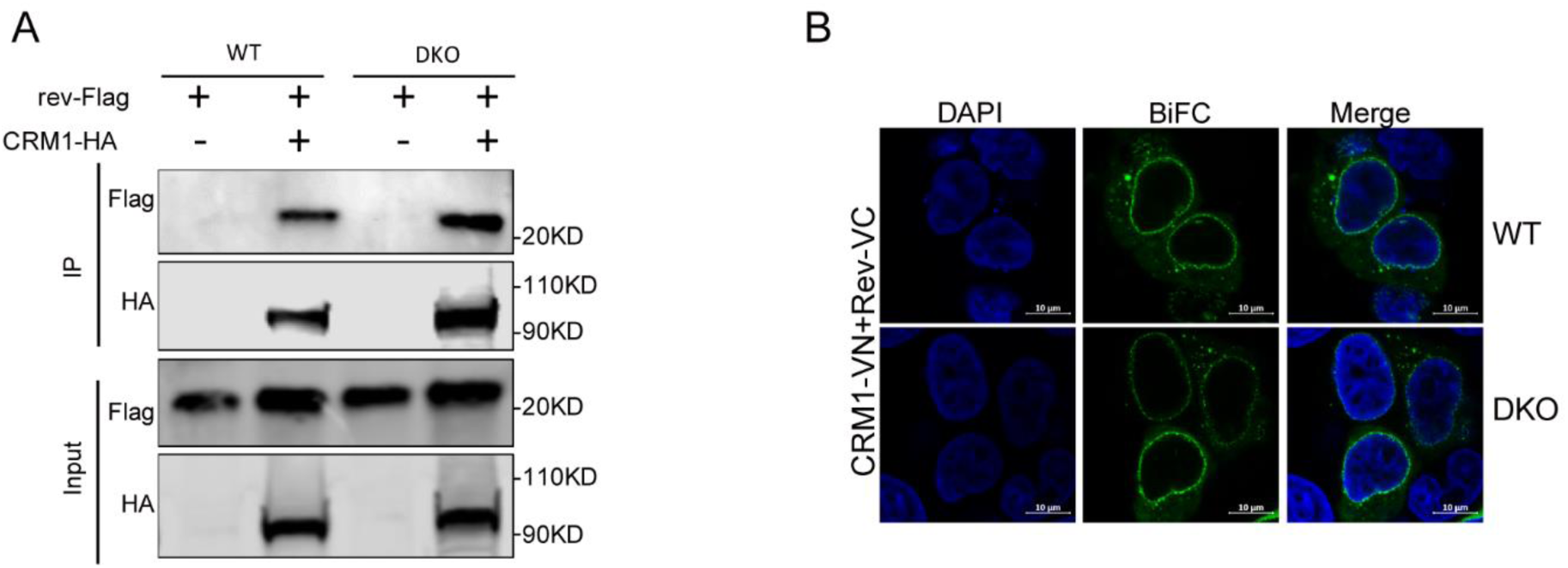
ANP32A/B not disrupt the interaction between Rev and CRM1 protein. (A) Detection of CRM1 interaction with Rev. 293T WT or DKO cells were co-transfected with plasmids expressing Rev-Flag along or in the presence of CRM1-HA vector, the whole cells lysates were prepared, immunoprecipitations were performed with anti-HA beads, and analyzed by Western blotting. (B) CRM1-VN and Rev-VC were co-transfected into 293T WT and DKO cells and the BiFC green fluorescent signals were visualized by confocal microscopy.

## DISCUSSION

Synthesis of viral structural components from full-length unspliced mRNA is a critical step in the production of viral progeny during HIV-1 replication. The HIV-1 Rev protein plays a key role in the nuclear export of unspliced and incompletely spliced viral RNA containing the RRE structure (45,46). The N-terminal domain of Rev binds directly with the RRE structure on the unspliced viral RNA, and the C-terminal domain of Rev binds to CRM1, therefore enabling the transportation of the RNA. Here we use a double knock out of ANP32A and ANP32B, generated using the CRISPR/Cas9 system, to provide novel evidence that the cellular factors ANP32A and ANP32B are both functional and essential components in the production of HIV-1. ANP32A and ANP32B bind to Rev and CRM1 and support the nuclear export of unspliced and partially spliced mRNA. Furthermore, our studies show that inhibition of the function of CRM1 or deletion of ANP32A and ANP32B, impaired unspliced viral RNA export and HIV-1 production. All these results demonstrated that ANP32A and ANP32B play an important role in HIV-1 production and presumably are key members of the Rev-CRM1 RNA export complex.

The evolutionarily conserved ANP32 family of proteins is characterized by an N-terminal variable number leucine rich repeat domain (LRR) and a C-terminal acidic tail (28,47). ANP32A and ANP32B share a highly conserved structure. Recent study has showed that chicken ANP32A is a major host factor affecting avian influenza viral polymerase activity in human cells (38); ANP32B can function as a target of the M protein of henipavirus to support virus replication (48). Moreover, ANP32A and ANP32B act as necessary cofactors in the export of nuclear RNA of foamy virus without a Rev-like protein (40). The CRISPR/Cas9 system and siRNA screen approaches enable us to achieve a clear background of certain protein express and have allowed the identification of various host co-factors that mediate HIV-1 replication (49,50). In this study, we found that double, but not single, deletion of ANP32A and ANP32B using the CRISPR/Cas9 system in an HEK293T cell line significantly decreases HIV-1 production and export of unspliced viral RNA. Reconstitution of ANP32A or ANP32B restores the gag expression. This result indicates that both ANP32A and ANP32B function similarly and contribute equally to the export of viral RNA.

Many host factors have been reported to involve in the Rev dependent viral RNA export. Rev interacts with transport receptor CRM1 and other host factors, including Ran-GTP, DDX3, RMB14, and Naf-1, to facilitate RNA export (11,22,51,52). Certain proteins have been identified that bind not only to Rev, but also to CRM1. Examples include the naf-1 protein, which can interact with CRM1 to enhance mRNA nuclear transport and HIV-1 virus production (52); RBM14, which interacts with CRM1 and Rev protein and supports Rev-mediated export of unspliced viral transcripts (19); and UPF1, which is required in the regulation of vRNA nuclear export and which shuttles between the nucleus and the cytoplasm to form a multimeric complex containing UPF1, DDX3, CRM1 and Rev protein(53). There are also some factors such as Sam68, matrin 3, and DDX1, that interact with Rev and enhance export of viral RNA without evidence of binding to CRM1. Sam68 expression is required for Rev function by directly regulating the CRM1-mediated Rev nuclear export pathway but Sam68 itself does not actively shuttle between the nucleus and the cytoplasm (18,54,55). The nuclear matrix protein matrin 3 is shown to interact with Rev protein and bind Rev/RRE-containing viral RNA to increase cytoplasmic expression of these viral RNAs (56,57). DDX1 is known to physically interact with both Rev and the RRE and it can act through RNA to promote HIV-1 Rev-RRE assembly (20,58). Our results showed that ANP32A/B can bind to Rev and CRM1. Without ANP32A and ANP32B, hardly any *gag* mRNA was detected in the cytoplasm. From these results, we propose a key role of ANP32A/B proteins in the Rev/RRE-CRM1 RNA export pathway (Fig. 7).

**FIG 7.**
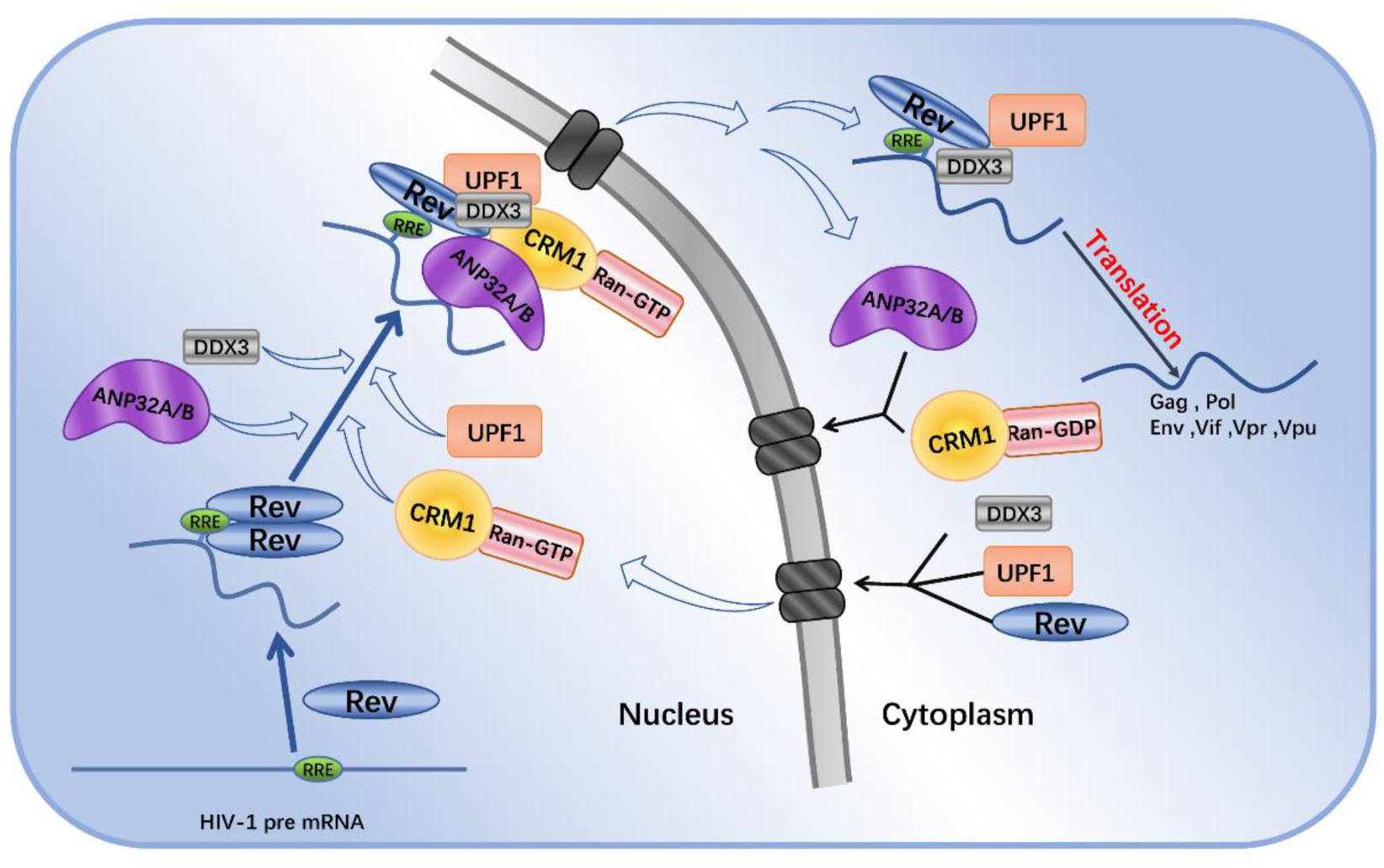
Proposed model for ANP32A/B function in CRM1-mediated nuclear export. In the late phase of HIV-1 replication, ANP32A/B interact with Rev/RRE and CRM1 complex and mediated unspliced mRNA nuclear transport via the nuclear pore complex. In the cytoplasm, the complex dissociated and several factors including ANP32A/B recycle back to the nuclear location. The unspliced mRNA translate to the viral protein or assemble virions as the viral RNA.

The mechanism of function of the Rev protein, and the details of transport of RNA need further investigation. In our work, using RNA Scope technology, we found that without either ANP32A or ANP32B, most of the *gag* mRNA is restricted to the nucleus. In addition, we evaluated the contribution of CRM1 and ANP32A/B in HIV-1 Rev dependent RNA transport. We found that blocking of CRM1 by LMB dramatically reduces gag expression in WT 293T cells, which is consistent with results from previous studies. Surprisingly, in DKO cells, HIV-1 has a low-level of virus production ability and inhibition of CRM1 by LMB showed only a slight reduction in viral replication (Fig. 5E). This result indicated that without ANP32A/B, the CRM1-mediated export pathway for HIV-1 RNAs was blocked. Thus, all our experiments appear to support the conclusion that either ANP32A or ANP32B is required to facilitate nuclear export of HIV-1 RNA though the Rev/CRM1 pathway.

We have observed that viral unspliced RNAs at 24h post infection was predominantly located in the nucleus in the DKO cell while most of it has transported to the cytoplasm in the WT cells. Over accumulation of the RNA in the nucleus presumably blocked the process of RNA synthesis. In our study, we found that total *gag* RNA is significantly lower compared with WT (Fig. 2C). Whether nuclear retention is accompanied of degradation or over splicing of the unspliced mRNA needs to be investigated.

In summary, we found that ANP32A and ANP32B are essential host factors supporting HIV-1 virus production. ANP32A and ANP32B specifically interact with Rev and CRM1 and both contribute to the export of HIV-1 unspliced RNA. The mechanism of how ANP32A and ANP32B function in the Rev/RRE-CRM1 complex needs further investigation.

## EXPERIMENTAL PROCEDURES

### Cell lines

Human embryonic kidney HEK293T and HeLa cells were maintained in Dulbecco’s modified Eagle’s medium (DMEM, Gibco) with 10% fetal bovine serum (FBS; Sigma), 1% penicillin and streptomycin (Gibco), and kept at 37℃ with 5% CO_2_. HEK293T cells were used to generate ANP32A and ANP32B gene knockout cells using CRISPR/Cas9 technology. Briefly, we designed the gRNA sequence from the gRNA design tool (http://crispr.mit.edu/) and inserted it into the pMD18-T backbone vector which contains the U6 promoter, gRNAs specific for host factors, a guide RNA scaffold, and a U6 termination signal sequence. Then the Cas9-eGFP expression plasmid (pMJ920) (provided by Jennifer Doudna) was co-transfected together with gRNA expression plasmids into HEK293T cells. GFP-positive cells were sorted by fluorescence-activated cell sorting (FACS) and cloned by limiting dilution. The positive clones were screened by Western blotting. Single knockout (KO) cell lines ∆ANP32A (AKO) and ∆ANP32B (BKO), and double KO cell line (DKO) were successfully generated. Stable cell lines expressing human ANP32A and human ANP32B in DKO cells, named DKO+A, DKO+B, were generated through retroviral transduction. Briefly, HEK293T cells in 6-well plates were transfected with 2.0 μg pLPCX, 2.0 μg pCGP and 0.25 μg VSV-G expression plasmids by Lipofectamine^®^ 2000 Transfection Reagent with recommended protocols as described previously(59). The pseudotype MLV virus expressing the ANP32 protein was harvested 48 h after transfection and used to infect the DKO cells. Positive cells were sorted using puromycin (1 μg/ml).

### Plasmids

The HIV-1 luciferase (Luc) reporter proviral vector pNL4-3-lucΔVifΔEnv was a gift from Dr Yong-Hui Zheng in Michigan State University. The pCAGGS plasmids containing human ANP32A and ANP32B were kindly provided by Dr. Wendy S. Barclay. The human CRM1 gene was cloned from cDNA and inserted into a pcDNA3.1 expression vector with 2×HA tag at the C terminus. The pcDNA3.1-Rev-HA plasmid was provided by Dr. Wang Jianhua from the Institute Pasteur of Shanghai, Chinese Academy of Sciences.

The plasmids of pCAGGS-ANP32A-VN-FLAG, pCAGGS-ANP32A-VC-FLAG, pcDNA3.1-HA-Rev-VC, pcDNA3.1-CRM1-VN-HA were constructed by fusing the N-terminal residues 2-173 of Venus (VN) and C-terminal residues 174-239 (VC) to the C-terminus of ANP32A, Rev or CRM1. The VN was inserted into pCAGGS-ANP32A-VN-FLAG and pcDNA3.1-CRM1-VN-HA between gene and HA or FLAG tags. The VC was inserted into the N-terminus of pcDNA3.1-FLAG-trim5α plasmids. While the pcDNA3.1-HA-Rev-VC was constructed by inserting the VC followed the rev gene sequence.

### VSV-G pseudotyped retrovirus production

Pseudotyped viruses of HIV-1 were produced by transfection of viral genome constructs in HEK293T cells as described previously(59). In brief, for rescue of the HIV-1 pseudovirions, HEK293T cells were seeded in 100-mm dishes and transfected with pNL4-3-lucΔVifΔEnv and vesicular stomatitis virus glycoprotein (VSV-G) using the PEI transfection reagent (Polysciences) following the manufacturer’s instruments. Forty-eight hours later, the culture supernatants were collected clarified by low-speed centrifugation and then were centrifuged at 25,000 rpm at 4 °C for 2 h using a Beckman SW-41Tirotor. Viral pellets were resuspended in lysis buffer and subjected to Western blotting analysis.

### Cell fraction and real-time PCR analysis

WT and DKO cells were seeded into 6-well plates and infected with HIV-1_NL4-3_ pseudotyped virus for 24 h. To measure viral reverse transcripts, virus stocks were first treated with 50 units/ml RQ1-DNase (Promega) for 1 h to remove any plasmid DNA. Both WT and DKO cells were incubated at 37 ℃ with these viruses. Cells were collected at 2, 6, and 18 h post infection and the total cellular DNA was extracted using DNeasy kits (Qiagen). Equal amounts of cellular DNA after a further treatment with *Dpn* Ⅰ were then subjected to real-time PCR in order to measure early and late viral reverse transcripts, as described previously (61). β-globin DNA was measured as an endogenous control. The primers used are as follows: early oHC64-F, 5-TAACTAGGGAACCCACTGC-3, early oHC64-R, 5-GCTAGAGATTTTCCACACTG-3; late RT MH531-F, 5-TGTGTGCCCGTCTGTTGTGT-3, late RT MH532-R, 5-GAGTCCTGCGTCGAGAGAGC-3; β-globin-F, 5-CCCTTGGACCCAGAGGTTCT-3, β-globin-R, 5-CGAGCACTTTCTTGCCATGA-3. To quantify the mRNA in the WT and DKO cells, total RNA from the cells was extracted using a Bio-fast simply P RNA extraction kit (catalog # BSC60S1, Bioer). Nuclear and cytoplasmic RNA fractions were purified with equal volume of elution buffer by using a PARIS Protein and RNA isolation kit (Ambion, Life Technologies) following the manufacturer’s instructions. Equal volume RNA was reverse transcribed into cDNA using the PrimeScript RT Reagent Kit with gDNA Eraser (Takara) according to the manufacturer’s instructions. Real-time PCR was performed using the SYBR green PCR mixture to calculate the absolute quantification with standard curves for unspliced RNA (*gag*) and completely spliced RNA (*tat*). The primers used are as follows: Gag-forward, 5-GTGTGGAAAATCTCTAGCAGTGG-3, Gag-reverse, 5-CGCTCTCGCACCCATCTC-3 (52); Tat-forward, 5-CAGCCTAAAACTGCTTGTAC-3, Tat-reverse, 5-GGAGGTGGGTTGCTTTGATA-3.

### Formaldehyde cross-linking, Co-immunoprecipitation and Western Blotting

To examine the interactions between proteins in cells, HEK293T cells were transfected with the indicated plasmids using the PEI transfection reagent. Immunoprecipitations were performed using an anti-HA affinity gel (catalog NO. A2095; Sigma-Aldrich) following the manufacturer’s instructions. Briefly, cells co-transfected with indicated plasmids were harvested after 48 h and washed with PBS. Formaldehyde cross-linking was subsequently performed. Firstly, cells were crosslinked with 1% formaldehyde for 20 min at RT and quenched by the addition of 0.25M glycine for 5 min. Then the cells were lysed with lysis buffer containing protease inhibitor cocktail (catalog NO. P8340; Sigma-Aldrich), sonicated and centrifuged at 13,000 g for 10 min. 30 μl of the anti-HA affinity gel was washed with ice-cold PBS and incubated with the cell lysates overnight at 4°C. Following the incubation, beads were washed with lysis buffer, and bound proteins were eluted by mixing and heating the beads in sample loading buffer for 5 min at 98°C. Samples were loaded on a 12% Tris gel (Bio-Rad). The gels were transferred to nitrocellulose membranes and blocked with 5% nonfat dry milk (NFDM) for 1 h and incubated with the appropriate primary antibodies. After extensive washing with TBST, the gels were further incubated with the relevant secondary antibody for 1 h at RT. Target protein bands were detected and analysis performed using Licor Odyssey (USA).

### Confocal imaging

HeLa cells were seeded onto 35-mm glass-bottomed culture dishes (Eppendorf) and then co-transfected with the indicated plasmids. Thirty-six hours later, the dishes were washed with ice-cold PBS (PH 7.6) and the cells were fixed with 4% para-formaldehyde for 30 min at RT. The dishes were then washed with PBS three times and subsequently permeabilized in 0.1% Triton X-100 solution for 15 min at RT. The cells were then treated with 5% BSA blocking solution for 1 h and the incubated with mouse monoclonal anti-HA (Sigma) antibody and rabbit anti-Flag antibody (Sigma) solution for 1 h at RT. The secondary mouse conjugated antibody to Alexa Fluor 488 and secondary rabbit conjugated antibody to Alexa Fluor 633 (Thermo) were subsequently added and incubated for a further 1 h, followed by washing with PBS three times. DAPI (Beyotime, China) staining was performed to visualize nuclei. Stained sections were captured and processed by fluorescence microscopy (ZEISS, LSM880, Germany).

### RNA in situ hybridization (RNAISH)

Locations of target mRNA were detected by ISH using the RNA-Scope Multiplex Fluorescent Reagent Kit v2 (Advanced Cell Diagnostics) according to the manufacturer’s instructions. In brief, HEK293T cells were cultured on cell imaging Dishes 35*10mm (Eppendorf) and transfected. Twenty-four hours later, the cells were fixed with 4% para-formaldehyde for 30 min and permeabilized for 15 min. The cells were then incubated in the HybEZ™ Oven for 2 h at 40 ℃ to complete the RNA hybridization step. The *gag-pol* mRNA probe (Advanced Cell Diagnostics, catalog # 317691) was designed and synthesized by Advanced Cell Diagnostics, and consisted of 20 pairs of oligonucleotides of the target mRNA transcript. Following proprietary preamplification and amplification steps, target mRNAs were labeled with certain fluorescence. DAPI staining was performed to visualize nuclei. Stained sections were captured and processed by fluorescence microscopy (ZEISS, LSM880, Germany).

### Statistical Analysis

All experiments were performed independently at least three times, with a representative experiment being shown. All statistical analyses were performed in the GraphPad Prism using student’s t test for pairwise and one-way analysis of variance (ANOVA) for multiple comparisons. *p < 0.05, **p < 0.01, ***p < 0.001, ****p < 0.0001, NS, not significant (p > 0.05).

## ACKNOWLEDGEMENTS

The authors wish to thank W. Barclay, Dr. Hui Zhang, Dr. Yong-Hui Zheng, and Dr. Jianhua Wang for provision of the plasmids.

## AUTHOR CONTRIBUTIONS

YW, XW designed experiments and wrote the manuscript; YW, HZ, LN, DC, ZZ performed the experiments; LN, YZ, XW analyzed the data. All authors reviewed the results and approved the final version of the manuscript.

